# Species-specific metabolic reprogramming in human and mouse microglia during inflammatory pathway induction

**DOI:** 10.1101/2023.03.10.531955

**Authors:** Angelica Maria Sabogal-Guaqueta, Alejandro Marmolejo-Garza, Marina Trombetta Lima, Asmaa Oun, Jasmijn Hunneman, Tingting Chen, Jari Koistinaho, Sarka Lehtonen, Arjan Kortholt, Barbara M. Bakker, Bart J.L. Eggen, Erik Boddeke, Amalia Dolga

## Abstract

Metabolic reprogramming is a hallmark of the immune cells in response to inflammatory stimuli. This metabolic process involves a switch from oxidative phosphorylation (OXPHOS) to glycolysis, or alterations in other metabolic pathways. However, most of the experimental findings have been acquired in murine immune cells and little is known about the metabolic reprogramming of human microglia. In this study, we investigated the transcriptomic and metabolic profiles of mouse and iPSC-derived human microglia challenged with the TLR4 agonist LPS. We found that both species displayed a metabolic shift and an overall increased glycolytic gene signature in response to LPS treatment. The metabolic reprogramming was characterized by the upregulation of hexokinases in mouse microglia and phosphofructokinases in human microglia. This study provides the first direct comparison of energy metabolism between mouse and human microglia, highlighting the species-specific pathways involved in immunometabolism and the importance of considering these differences in translational research.

## Introduction

Microglia are the resident innate immune cells of the central nervous system (CNS) and are involved in the immune response to pathogens or alteration to the CNS microenvironment. Microglia are also required for neurodevelopment, neuroplasticity, and tissue repair^1–3^. Dysregulation of microglial function has been well established in pathologies linked to neurodegeneration, including Alzheimer’s disease (AD)^4–6^, Parkinson’s disease^7^, multiple sclerosis,^8,9^ and Huntington’s disease^10^. The lack of disease-modifying therapies for these conditions demonstrates the need to better delineate mechanisms that govern microglial function and to study the modulation of key mediators of such mechanisms for potential therapies.

Microglia acquire an inflammatory phenotype comprising production of pro-inflammatory cytokines interleukin-1β (IL-1β) and tumor necrosis factor-α (TNF-α) in response to pathogenic stimuli. Besides the inflammatory phenotype, microglia can adopt several phenotypes to clear pathogenic substances, eliminate cellular/synaptic debris, clear metabolic waste, regulate synaptic pruning, neuronal maturation, and support neuro-regeneration. During phenotypic transition, metabolic reprogramming is considered a hallmark of inflammatory murine macrophages/microglia^11,12^. Metabolic reprogramming or a metabolic switch encompasses a variety of cellular alterations in bioenergetic pathways to adapt to cell’s metabolic needs. The changes in metabolic pathways include several processes, such as oxidative phosphorylation (OXPHOS), tricyclic acid cycle (TCA cycle), glycolysis, the pentose phosphate pathway, amino acid metabolism, and fatty acid oxidation. Recent studies have shown that mouse primary microglia are able to switch their cell metabolism from mainly mitochondrial OXPHOS to glycolysis^12^ in response to pro-inflammatory stimuli, such as lipopolysaccharides (LPS). Conversely, amyloid-β (Aβ) elicits differential immune responses^13^ in acute and chronic manners, pointing at a role of metabolism in trained innate immunity in microglia. Similarly, this phenomenon of metabolic switch to glycolysis is also present in mouse macrophages, dendritic cells, NK cells, B cells and effector T cells^14,15^.

Under physiological conditions, immune cells, including microglia and macrophages, primarily rely on oxidative phosphorylation^16^. Under inflammatory conditions, an accumulation of citrate and succinate has been reported in macrophages, which contributes to an increase in reactive oxygen species (ROS) and nitric oxide production, leading to a switch from anti- to pro-inflammatory phenotype in the immune cells^14,17^. These findings were followed by an exponential surge of interest in reprogramming metabolic pathways and the term of “immunometabolism” was coined in 2011 by Mathis & Schoelson^18^. However, the vast majority of these experimental studies have been performed in murine immune cells and not much information is available on metabolic reprogramming in human immune cells, specifically innate brain immune cells, microglia. For instance, species-specific differences in LPS-treated macrophages have been reported in mouse and human systems^19^ indicating that murine findings shall be interpreted with caution and highlighting the need to establish more adequate human model systems.

Studies of human brain microglia have been performed on isolated microglia from fresh post-mortem samples from potentially neuropathologically affected individuals, which might be hindered by a high interindividual variation. Alternately, robust differentiation protocols of iPSC-derived human microglia could provide a possibility to study metabolic profiles. First studies on iPSC-derived microglia were documented in 2016 and mainly focused on the characterization of the microglial phenotype in terms of differentiation and maturation. Hence, the goal of our study was to investigate the metabolic reprogramming in human iPSC-derived microglia compared with mouse microglial *in vitro* and *in vivo* models in response to the prototypical stimulus LPS. We observed dysregulation of metabolic pathways concomitant with upregulation of inflammatory pathways in both *in vivo* and *in vitro* treated mouse and human iPSC-derived microglia. Functional measurements demonstrated glycolytic upregulation in both species but discrepant changes in oxidative metabolism with a decrease in mouse but not in human microglia. This report provides evidence on species-specific differences in metabolic reprogramming in microglia.

## Results

### LPS-stimulation of mouse microglia induces an inflammatory gene signature

LPS is commonly used to investigate the stimulation of immune cells and their response to TLR4 activation, as a model to mimic inflammatory processes. To study how microglia respond to LPS at the transcriptome level, we performed RNA-seq of LPS-treated primary mouse microglia (Fig 1A). We analyzed the RNA-seq data following LPS challenge for a short period (4h) and a longer period (24h) of mouse microglia to assess acute and late responses to LPS stimulus. The analysis of these data revealed sets of LPS-induced genes (Table S1) and robust transcriptomic changes in mouse microglia (Fig 1B). LPS-treated microglia, regardless of the treatment duration segregated from control microglia in the first principal component. Interestingly, transcriptomic differences between LPS treatments for 4h and 24h segregated microglia in the second component (Fig 1B).

**Figure 1.**
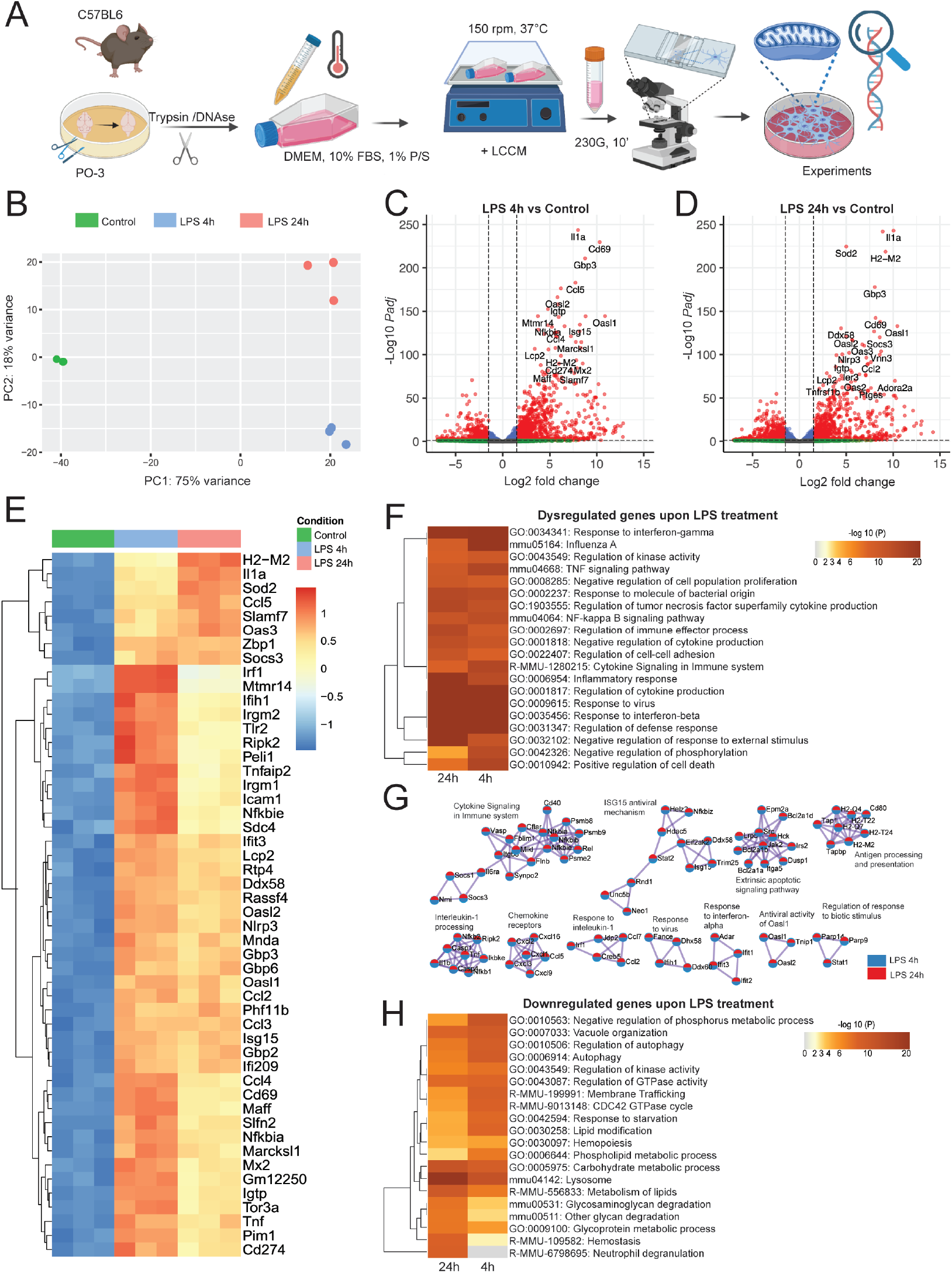
Transcriptomic analysis of LPS-treated primary mouse microglia. A. Experimental workflow of primary mouse microglia challenge. B. Principal Component Analysis (PCA) plot depicting each RNA-seq sample transcriptome, colored by experimental group. C - D. Volcano plots depicting fold changes and -Log2 of the adjusted p value per gene comparing 4 (C) or 24 (D) hr LPS treatment and the untreated control (24h). E. Heatmap depicting the top 50 most significant differentially expressed genes of 4hr LPS treatment compared with the control. Color key corresponds to row Z-score. F. Gene set enrichment analysis (GSEA) from the DEGs of LPS application of 4 and 24hr, depicted as a hierarchically clustered heatmap. Color key corresponds to the -Log10 of the p value for enrichment score. G. Protein-protein interaction network analysis: proteins from the lists of DEGs of cells challenged with LPS for 4 and 24hr, where Molecular Complex Detection (MCODE) algorithm was applied to identify densely connected network neighborhoods, where such neighborhood component is more likely to be associated with a particular complex or functional unit than the rest of the network. Legend denotes whether the components were found in the DEG lists of either comparison. H. GSEA from the downregulated genes in mouse microglia challenged with LPS for 4 and 24hr, depicted as a hierarchically clustered heatmap. Color key corresponds to the -Log10 of the p value for enrichment score. Part of this figure was created with BioRender.com.

Treatment with LPS for short time (4 h) induced a transcriptomic response mainly characterized by the upregulation of inflammation-related genes, such as IL1/chemokine signaling pathways (*Il-1a, Nfkbia, Ccl4, Ccl5*), ribonuclease activity (*Oasl2, Oasl1*), or immune cell response (*Cd69, Gbp3, Isg15, Igtp, Mtmr14, H2-M2*) among others. Short-term LPS treatment led to the downregulation of several genes related to immune cell differentiation, (such as *Inpp5d, Lhfpl2, Cebpa, Rab7b, Nfic, Kif21b, IL16ra, Tmem86a, Tmem104, Fblim1, S1pr1)* (Fig 1C) when compared to control cells. Treatment with LPS for longer time (24h) induced the expression of inflammatory genes *Il-1a, Nlrp3*, interferon-induced genes *Gbp3, Cd69, Oasl1*, immune activation genes *H2-M2, Ddx58, Socs3, Vnn3, Igtp*, and the mitochondrially-localized *Sod2* and decreased the expression of microglial markers *Pla2g15, Cx3cr1, Cd300lb* and of genes related to Alzheimer’s Disease such as *Pald1, Igf1, Plau, Flt1*, and the immune cell differentiation gene *Cebpa* (Fig 1D) when compared to the control cells. Hierarchical clustering of the top 50 (Fig 1E) most changed genes following 4h of LPS treatment showcased a module of upregulated genes, including many genes involved in the immune response, such as *Irf1, Mtmr14, Ifih1, Nfkbie, and Tnfaip2*. This cluster was downregulated following 24h compared to 4h LPS treatment. Another cluster of genes involved in the immune response, including *H2-M2, IL1a, Sod2, Ccl5* and *Socs3* was upregulated at a later stage after 24h LPS challenge.

Gene set enrichment analysis (GSEA) of statistically significant dysregulated genes following LPS application for 4 and 24hr demonstrated shared enrichment of inflammatory pathways such as NF-κB signaling pathway, inflammatory response, and TNF signaling pathway, among others (Fig 1F). We next applied Molecular Complex Detection (MCODE) algorithm on the Protein-Protein interaction (PPi) analysis that resulted from the input of DEGs of LPS-treated cells. This analysis resulted in a network characterized by top enriched clusters such as cytokine signaling in immune system, ISG15 antiviral mechanism, extrinsic apoptotic signaling pathway, antigen processing and presentation, interleukin-1 processing, among others (Fig 1G), depicting important cluster overlaps in the top enriched pathways both at 4h and 24h. Because the majority of the DEGs with largest fold changes were positively regulated upon LPS, we aimed to investigate the negative regulatory effect of LPS on gene expression. Accordingly, we performed GSEA of downregulated genes in microglia challenged with LPS and observed the highest shared enrichment in processes that comprise lysosomal metabolism (*Abca2, Lamp1, Cd63, Cd68, Hexa, Npl*), carbohydrate catabolism linked to glycolysis (*Pfkfb2, Akt1, Gpi1, Slc2a8*), pentose phosphate pathway (*H6pd*) and TCA cycle (*Mdh1*), and lipid catabolism (*Abcd1, Cpt1a, Gpx4, Acox3*), fatty acid synthesis (*Fasn*), autophagy (*Ulk1, Atg13, Atg7, Ubqln2*) and response to starvation (*Pparg, Apoe, Cd68, Foxo1, Foxo3, Jun*) (Fig 1H). Collectively, these results strongly suggest that upon LPS challenge, irrespective of the treatment duration, mouse microglia upregulate a battery of genes that are related to inflammation and immune activation, which is paralleled by downregulation of catabolic processes that aim to provide energy from substrates such as lipids, fatty acids, and cell organelles.

### LPS challenge of mouse microglia *in vitro* and *in vivo* induces transcriptomic signatures consistent with glycolytic upregulation

Our observations indicated that catabolic pathways such as lysosomal lipid degradation, pentose phosphate pathways and autophagy were downregulated in *in vitro* LPS-treated microglia. Based on these observations, we hypothesized that the metabolic reprogramming induced by LPS in microglia preferentially involve a shift towards glycolysis as an upregulated metabolic pathway to support immune activation. In-depth analysis of the gene expression involved in glycolysis demonstrated an upregulation of *Hk2, Hk3, and Pfkp*, and a downregulation of *Gapdh, Pfkfb2* and *Pfkfb4* following 4h of LPS challenge. Interestingly, *Hk2* continued to be upregulated following 24h of LPS treatment, while *Pfkfb3, Pfkfb4, Aldh2, Ldhb, Aldh9a1*, and *Gpi1* were downregulated (Fig 2A).

**Figure 2.**
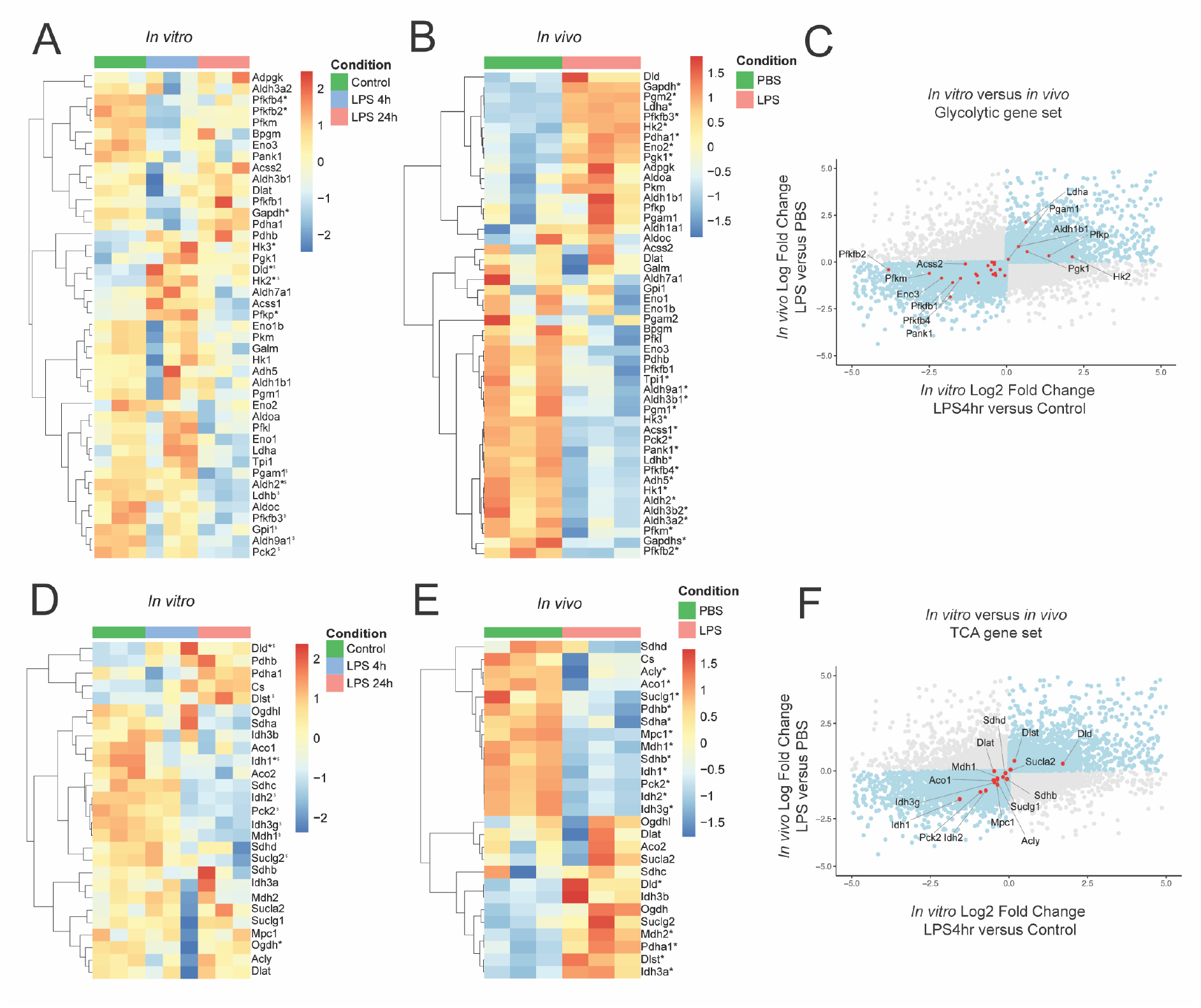
Transcriptomic changes in the glycolytic and TCA pathways in LPS-treated primary mouse microglia and *in vivo* LPS-treated mice. A,B. Hierarchically clustered heatmaps depicting gene expression changes in glycolytic pathways in in vitro cultivated mouse microglia (A) and in isolated microglia from LPS-treated mice (depicted as in vivo) (B). C. Scatterplots that graphically represent the fold changes of genes following LPS stimulation in vitro versus in vivo. Dots in blue depict genes that are increased or decreased in both models (in vitro and in vivo). Dots in red represent glycolytic genes. Color key corresponds to row Z-score. D,E. Hierarchically clustered heatmaps depicting gene expression changes of TCA in vitro (D) and in vivo (E). Color key corresponds to row Z-score. F. Scatterplots that graphically represent the fold changes of LPS stimulation in vitro versus in vivo. Dots in blue depict genes that are increased or decreased in both models. Dots in red represent TCA genes.

To assess whether the LPS-induced transcriptomic changes in cultured microglia reflect the in vivo microglia response to LPS, we compared the gene expression profiles of LPS-treated cultured microglia with microglia isolated from mice, 3 hr after an i.p. injection with LPS ^20^, with a particular focus on genes of the glycolytic pathway (Fig 2B). The analysis of glycolysis-related genes showed consistent downregulation of *Pfkfb2, Pfkfb1, Pfkfb4, Eno3* and consistent upregulation of *Hk2* and *Ldha* both *in vivo* and *in vitro* (Fig 2C). Hexokinases are critical enzymes in glycolysis as they phosphorylate glucose and initiate glycolysis. Interestingly, protein products of both isoenzymes can be detected in primary mouse-cultured microglia. *Hk3* was downregulated *in vivo* but not *in vitro*. On the other hand, *Pfkfb3* was upregulated *in vivo* but downregulated *in vitro*.

Glucose transporters (GLUTs) are key players in glucose metabolism and are not classically contained in the glycolytic gene sets (e.g. Gene Ontology or Kyoto Encyclopedia of Genes and Genomes). We hypothesized that GLUTs expression would be enhanced by LPS treatment in mouse microglia. Interestingly, we observed an upregulation of the transcripts that code for GLUT6 and GLUT8 (*Slc2a6*, and *Slc2a8*, respectively) upon LPS 4h and 24h treatment, and GLUT1 (*Slc2a1*) only at 4h after LPS treatment.

All in all, mouse microglia *in vitro* and *in vivo* exhibit shared dysregulation of glycolysis-related and other central metabolic gene transcription following LPS challenge consistent with glycolytic upregulation by increased GLUTs and HKs expression.

### LPS challenge of mouse microglia *in vitro* and *in vivo* induces transcriptomic signatures consistent with TCA dysregulation

Immune activation in macrophages and microglia has been linked to metabolic changes and adaptation in glycolysis coupled to the TCA cycle^13,19,21^. The final product of glycolysis is pyruvate, which can give rise to either lactate via lactate dehydrogenase (LDH) or to acetyl-CoA via pyruvate dehydrogenase (PDH). Acetyl-CoA enters the tricarboxylic acid (TCA) cycle generating NADH and FADH, which can subsequently fuel the electron transport chain. We hypothesized that the TCA cycle and PDH would be downregulated coupled to an increase of glycolysis to promote preferential glycolysis and aerobic fermentation into lactate. Indeed, we observed downregulation of specific genes involved in TCA cycle such as *Idh1* and *Ogdh1*, but upregulation of *Dld* following 4hr of LPS treatment. 24hr of LPS treatment downregulated *Idh1, Mdh1, Idh2, Idh3g, Suclg2* and upregulated *Dld* and *Dlst* (Fig 2D).

Analysis of *in vivo* LPS-treated microglia showed downregulation of TCA cycle genes *Idh1, Mdh1, Sdhb, Idh2, Pck2, Aco1*, among others and upregulation of *Dld, Suclg2, Dlst, Idh3a, and Idh3b* (Fig 2E). When comparing the transcriptomic changes of microglia upon LPS treatment *in vivo* and *in vitro*, we observed that the strongest up- and downregulated shared genes of the PDH complex and the TCA cycle were Dihydrolipoamide dehydrogenase (*Dld*, upregulated) and Isocitrate dehydrogenase (*Idh*, downregulated), respectively, (Fig 2F). *Idh* is the first of four oxidative steps within the TCA cycle and is the key step of this cycle. *Dld* codes for a mitochondrially-localized protein which serves as a NAD+ oxidoreductase in the pyruvate dehydrogenase multienzyme complex (*Pdhc*; *Pdc*; *Pdh*). This complex is an associated set of three enzymes that ultimately converts pyruvate to acetyl coenzyme A (acetyl-CoA). Collectively, changes in the expression of the TCA genes strongly indicate a decrease in TCA cycle activity following LPS challenge, indicating that in mouse microglia, TCA genes are regulated during the early (4h) inflammatory pathways.

### LPS challenge of human microglia induces an inflammatory gene signature

It has been proposed previously that mouse and human macrophage responses to TLR4 agonism could lead to differential outcomes in terms of metabolic reprogramming^11^. To further investigate whether this phenomenon occurs in microglia, as well, we generated iPSC-derived microglia-like (iMGL) cells, challenged them with LPS, and assessed their gene expression with RNA-seq profiling. We generated iMGL cells from control iPSCs and differentiated them by using a modified method described initially by Fatorelli and colleagues^22^. Microglial progenitors were further differentiated with IL-34, TFG-β, GM-CSF, CD200, and CX3CR1 to assure a full maturation profile. Following 14 days of maturation, human iMGL cells expressed microglial specific markers, TMEM119 and IBA1. In addition, qPCR indicated upregulation of *P2YR12, TMEM119, CD11c* and low expression of *Nanog* and *NeuN*, markers for pluripotency and neurons, respectively, demonstrating the features and markers of mature microglia (Fig 3A). Differentiated human iMGL cells exhibited ramified morphologies with a small round cell body and possessed dynamic surveying movements. To further assess whether the gene profile of the differentiated human iMGL cells resembles the adult or fetal human brain microglia, we compared their transcriptome and demonstrated that the differentiated iMGL cells were similar to fetal/adult brain microglia^23,24^ and iMGL cells^25^ generated with other available protocols. Additionally, the transcriptome of iPSC-derived microglia clustered apart from CD14^+^/CD16^-^ and CD14^-^/CD16^+^ monocytes and dendritic cells (Fig 3B). We observed that in the microglial differentiation protocol, there was a substantial collection-dependent effect on the transcriptomes of the iMGL cells (Fig 3C). Accordingly, the collection number was included as a covariate in our analysis. However, it was evident that LPS elicited robust transcriptomic changes in these iMGL cells with non-challenged control, LPS 4h and LPS 24h challenged cells clustering separately.

**Figure 3.**
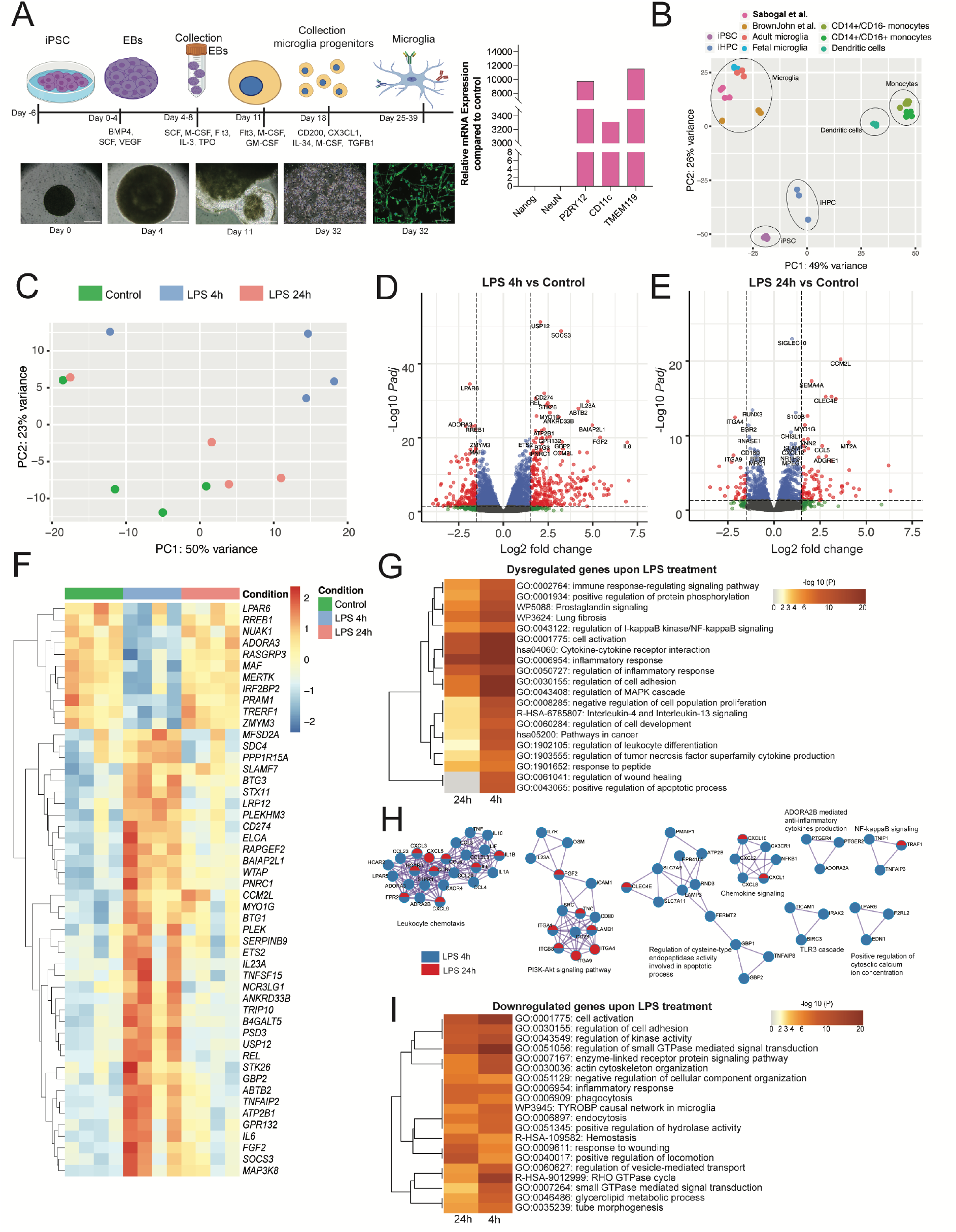
Transcriptomic analysis of LPS-challenged human iPSC-derived microglia. A. Differentiation protocol to generate iMGLs from human iPSCs. B. Principal Component Analysis (PCA) depicting transcriptomes of various cell types and previously published protocols for iPSC-derived microglia. C. PCA of untreated iPSC-derived microglia and LPS-stimulated microglia for 4 and 24hr. D,E. Volcano plots depicting fold changes and Log2 of the adjusted p value per gene by comparing 4 (D) or 24 (E) hr of LPS challenge and the untreated control (24h). F. Heatmap depicting the top 50 most significant differentially expressed genes of 4hr LPS treatment compared with the control cells. Color key corresponds to raw Z-score. G. Gene set enrichment analysis (GSEA) from the DEGs upon LPS 4 and 24hr depicted as a hierarchically clustered heatmap. Color key corresponds to the -Log10 of the p value for enrichment score. H. Protein-protein interaction network analysis: proteins from the lists of DEGs of cells challenged with LPS for 4 and 24hr, where Molecular Complex Detection (MCODE) algorithm was applied to identify densely connected network neighborhoods, where such neighborhood component is more likely to be associated with a particular complex or functional unit than the rest of the network. Legend denotes whether the components were found in the DEG lists of either comparison. I. GSEA of the downregulated genes following LPS 4 and 24hr challenge depicted as a hierarchically clustered heatmap. Color key corresponds to the -Log10 of the p value for enrichment. Part of this figure was created with BioRender.com.

Comparing the effects of LPS at different time points following LPS challenge, we observed that LPS-induced genes *USP12, SOCS3, IL23A, CD274, FGF2, CCM2L, IL23A, IL6* were strongly upregulated after 4h. *USP12* codes for a deubiquitinating enzyme, *SOCS3* codes for a protein member of the STAT-induced STAT inhibitor (SSI). *LPAR6, ADORA3, RREB1*, were downregulated after 4h following LPS challenge (Fig 3D). *LPAR6* codes for a Purinergic receptor from the P2Y family, *ADORA3* codes for the Adenosine receptor A3, *RREB1* codes for a zinc finger transcription factor that binds to RAS-responsive elements (RREs). After 24h LPS, *CCM2L, SEMA4A, CLECE, CCL5*, among others, were upregulated (Fig 2E). *CCM2L* codes for the Cerebral Cavernous Malformations 2 Protein-like, which has been reported to play a role in cell-adhesion and positive regulation of fibroblast growth factor (FGF) production, *CCL5* codes for a the chemoattract protein chemokine 5 ligand, *SEMA4A* codes for a member of the semaphoring family that has been implicated in nervous system development, *CLEC4E* codes for a member of the lectin-like superfamily, which are downstream targets of C-binding protein signaling. Genes such as *ITGA4* and *ITGA9* were downregulated upon LPS 24h. *ITGA4* and *ITGA9* code for protein members of the integrins, proteins that mediate cell surface adhesion. We identified a cluster of upregulated genes after 4h, but not after 24h, such as *IL6, CCM2L, DD274, FGF2, MAP3K8, REL, TNFAIP2, SLAMF7, IL23A, SOCS3* (Fig 3F). *IL6, IL23A, REL* code for key mediators in inflammatory responses. This battery of genes normalizes their gene expression in iMGL cells after 24h.

65 genes were dysregulated 24h after LPS and not after acute treatment with LPS for 4h. This list of genes contains *STEAP1B, IGFBP4, FYN, TREML3P, PLD4, SLC6A7, PTGES, FAXDC2, ITGA9, CCL7, IGF1, ENOX1*, among others. IFBP4 codes for a protein member of the insulin-like growth factor binding protein (IGFBP) family of proteins, which prolong the half-life of the IGFs. Interestingly, *IGF1* was upregulated at this timepoint. *SLC6A7* codes for a brain-specific L-proline transporter. *TREML3P* codes for a pseudogene in the TREM locus that have been suggested to play roles in TREM-dependent responses^26^. Notably, *ITGA9* codes for an alpha integrin, crucial for cell-cell and cell-matrix adhesion.

Similar to the mouse microglia, GSEA of statistically significant differentially expressed genes following LPS challenge of human iMGL cells demonstrated shared enrichment of inflammatory pathways such as NF-κB signaling pathway, prostaglandin signaling, and immune response (Fig 3G). For the iMGL cells, the number of DEGs after 24h was considerably lower than after 4h LPS treatment. To further illustrate this, we investigated the potential functional implications on the cells by predicting protein-protein interaction (PPi) maps from the DEGs for each comparison. Accordingly, we employed the MCODE algorithm on the PPi analysis resulting in a network characterized by top enriched clusters such as leukocyte chemotaxis, PI3K-Akt signaling, regulation of cysteine-type endopeptidase activity involved in apoptotic process, chemokine signaling, NF-kB signaling and positive regulation of cytosolic calcium ion concentration. This analysis demonstrated incomplete cluster overlaps in the top enriched pathways both at 4h and 24h, being 24h with the lowest enrichment. Data showcased a substantial decrease of the potential PPis after 24h, suggesting that the upregulated inflammatory pathways are dampened after 24h of challenge compared to short-term exposure (4h) (Fig 3H).

Next, we investigated the effects of LPS on human microglia gene expression of downregulated genes by performing GSEA. Interestingly, we observed a marked shared enrichment of the biological process regulation of small GTPase-mediated signal transduction (*CSF1, FBP1, IGF1, P2YR8, NOTCH1*), regulation of kinase activity (*CD86, MAP3K5, MAP3K5, ABCA7, CD4, CD74, TGFB1, TLR3*), glycerolipid metabolic process (*ABCD1, HEXA, HEXB, FAXDC2, ACSS2)*, and the Tyrobp causal network in microglia (*RUNX3, CD4, GPX1, ITGAM, LYL1, MAF, RGS1, TGFBR1, TYROBP, LOXL3*) (Fig 3I). Taken together, these data indicate that human microglia derived from iPSCs presented classical LPS-related transcriptomic responses coupled to downregulations of lipid synthesis, catabolism, and phosphate-based metabolism.

### LPS dysregulates the transcriptomic signature of glycolytic and TCA pathways in human iMGL cells

To follow-up on the transcriptomic dysregulations that LPS elicits on human iMGL cells, we hypothesized that the genes that code for enzymes of glycolysis would be dysregulated. We surveyed genes that code for components of this pathway and observed an upregulation of *ALDH1B1* and *PFKFB3*, and a downregulation of *PFKFB2, PFKFB4, ALDH3A2, HK1*, and *PFKM* at 4 hours compared to the control. Conversely, we observed upregulation of *PFKFB3, PKM, TPI1*, and *ENO3*, and a downregulation of gluconeogenesis genes (*FBP1)*, or other genes (*GALM, ALDH7A1, ALDH3A2)* at 24h post-LPS. These data strongly suggest that glycolytic flux of LPS-treated iMGLs may increase due to increased glucose uptake via GLUT transporters 3 and 6, with upregulation of *PFKFB3*.

For the TCA gene set, we observed a shared upregulation of ACO1 and downregulation of IDH1 and IDH2 at 4h and 24h LPS. These data highlight the upregulation of the genes whose protein expression would increase the TCA flux at 4h and 24h, with a downregulation of IDH.

### Glucose metabolism is differentially altered in mouse and human microglia challenged with LPS

To investigate how transcriptomic signatures are dysregulated across species upon the TLR4 agonist LPS, we analyzed one-to-one orthologues between mouse and human genomes and compared their fold-changes following LPS treatment. Our analysis demonstrated a shared downregulation of *PFKFB4* and *PFKM* and an upregulation of *DLD* in the *in vitro* cultured microglia following 4h of LPS challenge (Fig 4A). This gene signature (*PFKFB4* and *PFKM*) is characteristic for the glycolytic pathway. Interestingly, we observed differential induction of hexokinase isozymes: mouse microglia upregulated both *Hk2* and *Hk3*, while in human microglia *HK1* was downregulated and *HK3* had a tendency towards downregulation after 4hr LPS treatment (Fig 4B-D).

**Figure 4.**
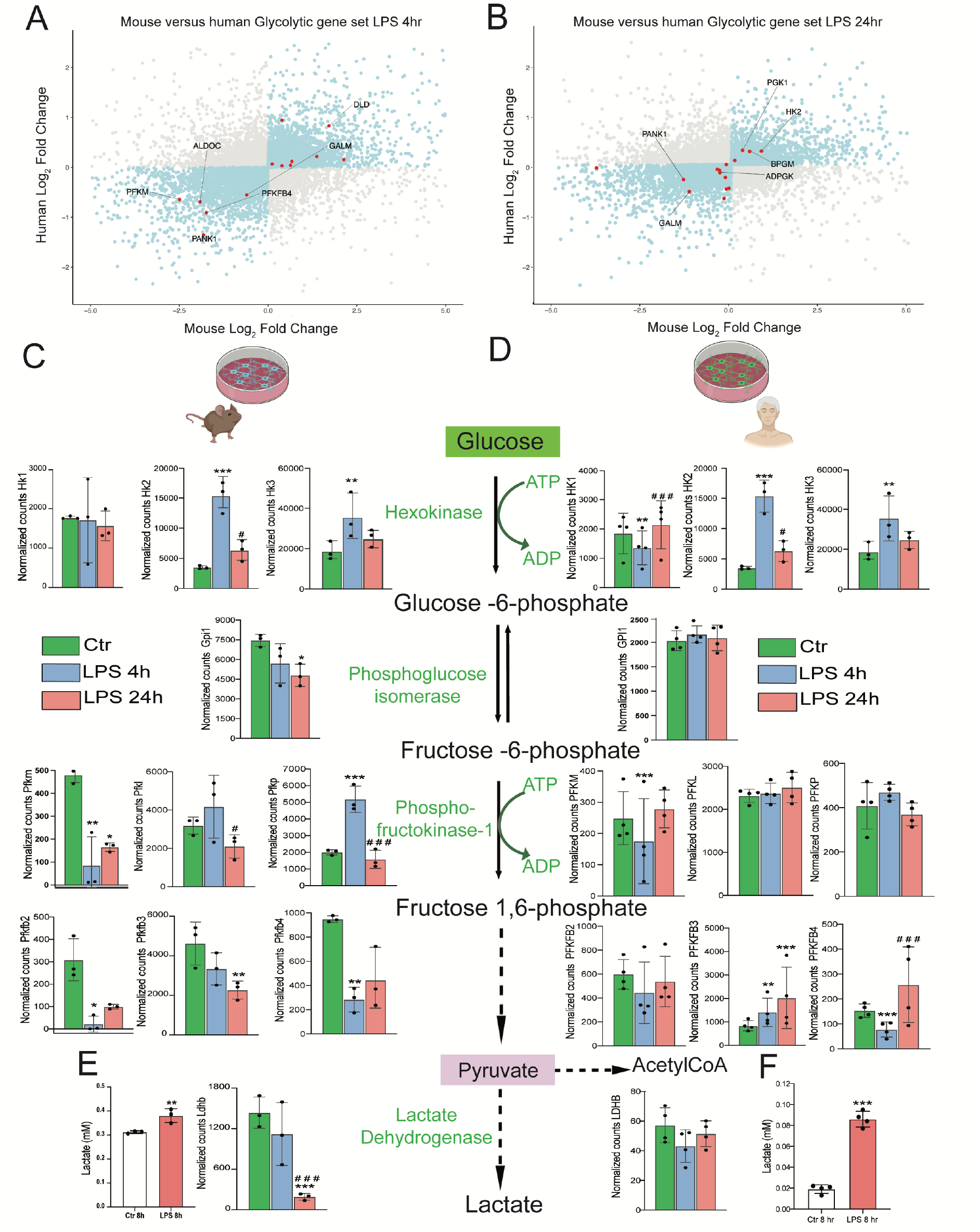
Transcriptomic comparison of mouse and human glycolytic genes. Scatterplots that graphically represent the fold changes of LPS stimulation in mouse and human microglia. Dots in blue depict genes that are increased or decreased in both models. In A and B, dots in red represent glycolytic genes at 4 and 24hr, respectively. In C we observed several of the main glycolytic genes regulated in mouse and in human (D) in the glycolytic pathway. E. Lactate release following treatment with LPS (250ng/ml) for 8 h in mouse and (F) human microglia (15,000 cells/well). Dotted lines denote intermediate steps in the pathway. Data are presented as mean ± SD, n = 3-4 per condition. *p < 0.05, **p < 0.01, ***p < 0.001 (compared with ctr), ^#^p < 0.05, ^##^p < 0.01, ^####^p < 0.001 (LPS 4h vs LPS 24hr). Part of this figure was created with BioRender.com.

After 24h, mouse microglia continued upregulating Hk2 and downregulating *Gpl1, Aldoc, Pfkl, Pfkb, Pfkfb3, Pfkfb4 and Ldhb*, while human microglia did not exhibit the same pattern (Fig 4B). *PFKFB3* was upregulated at 4h and 24h and *PFKB4* and *PFKM* were downregulated after 4h and no other isoform of PFKFB was dysregulated (Fig 4D) in the human microglia. The upregulation of hexokinase transcripts in mouse microglia suggests that there may be an increased glycolytic flux. Additionally, human microglia upregulated *PFKFB3*, an important regulator of glycolysis, since increased PFKFB3 activity increases the rate of glycolysis. Lactate dehydrogenases are enzymes that catalyze the reversible conversion of pyruvate and NADH to lactate and NAD^+^. To assess whether changes in gene expression levels are translated into functional data, we assessed lactate release as another readout of the glycolytic activity. The lactate determination using enzymatic lactate assay showed that after short term of LPS challenge (8h), lactate levels were increased in response to LPS in both mouse and human microglia compared to controls (Fig 4E, F). These data suggest that functional glycolytic microglial responses to LPS are conserved across species, but its key mediators may be different.

### Main enzymes from TCA cycle are regulated similarly in mouse and human microglia after LPS challenge

The final product of glycolysis is pyruvate, which can be converted to lactate or acetyl-CoA. In fact, the enzyme Dihydrolipoamide dehydrogenase (DLD) is one enzymatic component of the mitochondrial-based pyruvate dehydrogenase multienzyme complex in charge of the conversion of pyruvate to acetyl coenzyme A. We observed in human and mouse microglia an increase of DLD after 4h, followed by a reduction after 24h, as shown in the scatterplots and in the representative genes of the TCA pathway (Fig 5A-D). At the same time, the isocitrate dehydrogenase enzymes (IDH1 and IDH2) that catalyze the oxidative decarboxylation of citrate, resulting in 2-oxoglutarate, were downregulated in both species after 4h and continued to be downregulated after 24h, especially in human microglia (Fig 5D). Likewise, Succinyl-CoA ligase [GDP-forming] subunit beta enzyme, encoding for the *SUCLG2* gene, catalyzes the reversible conversion of Succinyl-CoA to succinate and acetoacetyl CoA was downregulated after 24h in both species, as shown in the scatterplot comparison and bar graphs (Fig 5B-D). Altogether our data show key enzymes such as *IDH, DLD, SUCLG2* are altered and their regulation might impact metabolic fluxes in response to LPS-mediated immune activation, thus supporting the metabolic reprogramming associated pathways.

**Figure 5.**
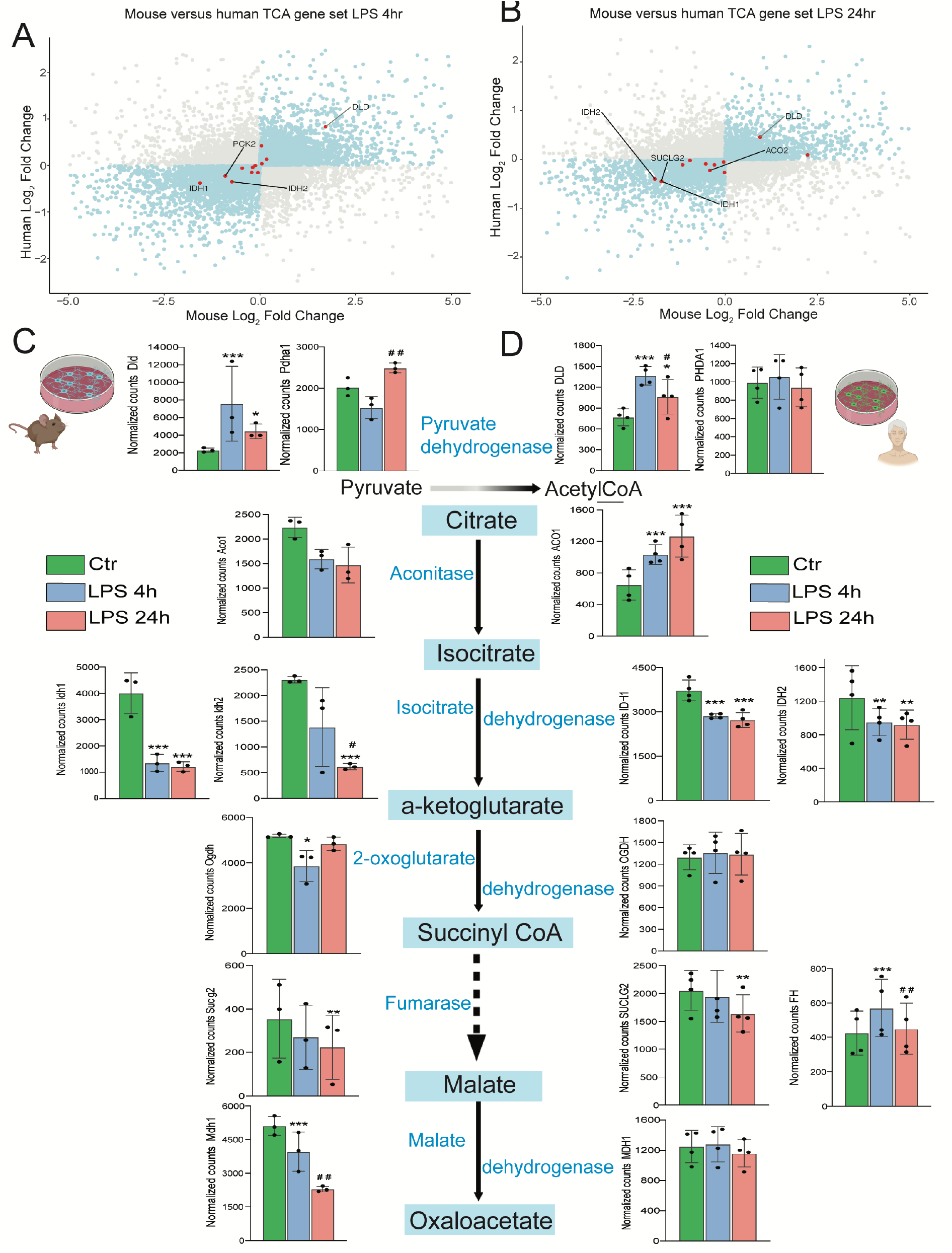
Transcriptomic comparison of mouse and human TCA cycle genes. Scatterplots represent the fold changes of LPS stimulation in mouse and human microglia. Dots in blue depict genes that are increased or decreased in both models. In A and B, dots in red represent TCA genes at 4 and 24hr, respectively. C-D. Several of the main TCA cycle genes are regulated in mouse (C) and in human (D) microglia. Dotted lines denote intermediate steps in the pathway. Data are presented as mean ± SD, n = 3-4 per condition. *p < 0.05, **p < 0.01, ***p < 0.001 (compared with ctr), ^#^p < 0.05, ^##^p < 0.01 (LPS 4h vs LPS 24hr). Part of this figure was created with BioRender.com.

### Species differences in metabolic reprogramming

To study the functional metabolic alterations in human vs mouse microglia, the bioenergetic profile of unstimulated and LPS-challenged microglia was evaluated using a Seahorse XF-analyzer. For functional analysis we included iMGLs derived from another iPSC line that was purchased from Gibco. The analysis of Oxygen consumption rate (OCR) data of mouse microglia showed that LPS does not affect mitochondrial respiration after a short challenge (4h), while long treatment (24h) decreased mitochondrial respiration compared to the control mouse microglia (Fig 6A). In contrast, the OCR of LPS-treated human microglia is decreased after 4h and increased after 24h compared to the unstimulated control (Fig 6B). We observed a decrease in the basal respiration of mouse microglia after 24h LPS treatment, however, we did not observe significant changes in the measurements of ATP-linked OCR and maximal OCR compared to the control (Figure 6C-E).

**Figure 6.**
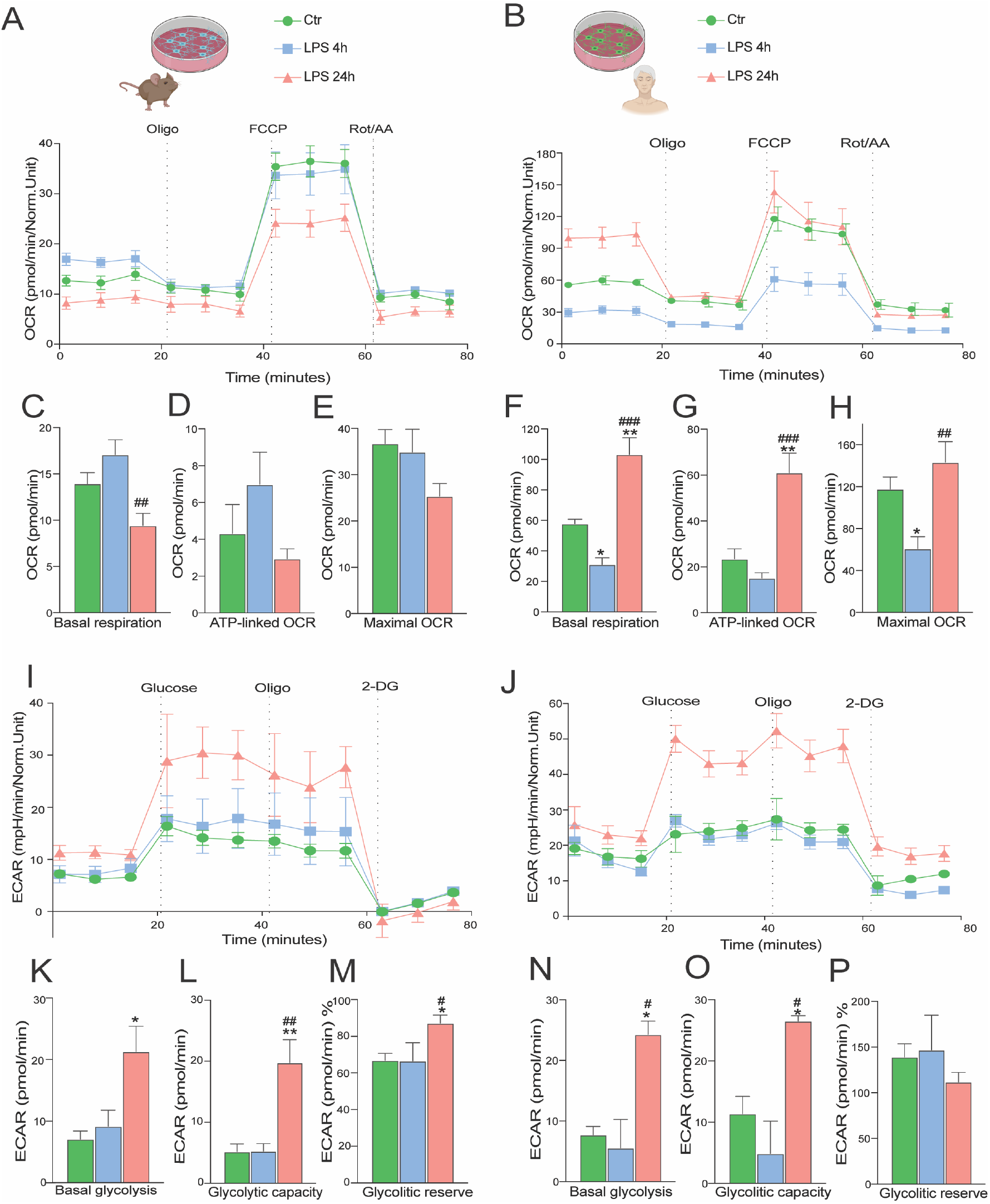
Mitochondrial respiration in mouse and human microglia. Representative experiment showing OCR (oxygen consumption rate) in A. mouse and B human microglial cells treated with LPS for 4 and 24hr. Measurements were obtained by an extracellular flux analyzer (Seahorse Bioscience). C-E. Bar graph showing different mitochondrial respiration parameters represented as mean ± SEM of the OCR values in mouse and F-H in human microglia. Representative experiment showing ECAR (Extracellular acidification rate) in I. mouse and J. human microglia cells treated with LPS for 4 and 24hr. K-M. Bar graph showing different mitochondrial respiration parameters represented as mean ± SEM of the ECAR values in mouse and N-P in human. Data are shown as mean ± SEM, n = 3–6 technical replicates. All experiments were repeated at least three times for each cell line (Ctr 8.2 & an iPSC line purchased from Gibco). p-values indicating statistically significant differences between the mean values are defined as follows: *p < 0.05, **p < 0.01 (compared with ctr), ^#^p < 0.05, ^##^p < 0.01(LPS 4h vs LPS 24hr). Part of this figure was created with BioRender.com.

Human microglia showed significance increase in basal respiration and ATP-linked OCR following 24h LPS treatment compared to the control (Fig 6F-G). Interestingly, the basal respiration and maximal OCR was decreased after 4h LPS treatment in human microglia (Fig 6F, H), while in the mouse microglia it was reduced (Fig 6C, E). On the other hand, the extracellular acidification rate (ECAR), an index of glycolysis feeding into lactate, was significantly increased after 24h LPS stimulation in both species (Fig 6I-J).

Basal glycolysis and glycolytic capacity were increased after 24h LPS stimulation in mouse (Fig 6K-L) and human microglia (Fig 6N-O), however, the glycolytic reserve was increased just in mouse microglia (Fig 6M, P). This indicates that the capability of microglia to respond to an energetic demand, as well as, how close the glycolytic function is to their maximum capacity depends on the species. These data indicate that LPS stimulation affects mitochondrial respiration in both mouse and human cells, and the overall response in basal and glycolytic capacity is similar between the two species. Importantly, the metabolic reprogramming towards an increased glycolytic activity was demonstrated in both mouse and human microglia.

## Discussion

In this study, we report that mouse and human microglia undergo metabolic reprogramming upon LPS stimulation. Overall, LPS treatment promoted glycolytic metabolism in both species. Interestingly, the increases in glycolytic activity are mainly attributed to hexokinase expression in mouse, and phosphofructokinase expression in human microglia. Both species upregulated glucose transporters. However, GLUT6 was only upregulated in human iMGLs at 4h LPS, while in mouse microglia continued to be upregulated until 24h.

Additionally, oxidative metabolism was suggested to be downregulated in both species by transcriptomic analysis, while functional experiments demonstrated a differential time profile of maximal FCCP stimulated flux: in mouse a slow decrease that becomes evident after 24hr, whereas in human microglia there is a fast, transient decrease (4hr) which is normalized after 24hr. This is the first report of cross-species transcriptional and functional responses of microglia challenged by inflammatory stimuli (Fig 7). We compared the transcriptomic response of mouse microglia in vitro and in vivo, since in vivo mouse microglia might behave differently compared to cultured acutely-isolated microglia^27,28.^ Overall, the induction of the glycolytic genes *Hk2, Pfkp* and *Pkm* in the *in vivo*-treated microglia was robust, and *Hk2* and *Pfkp* induction was recapitulated *in vitro* cultured cells, as well. Both models did not show significant dysregulation of *ldha* while a decrease of *ldhb* was detected only *in vitro* cultured cells. LDHA has a higher affinity for pyruvate, and depending on the metabolite concentrations preferentially converts pyruvate to lactate, and NADH to NAD+ under anaerobic conditions, whereas LDHB possess a higher affinity for lactate, preferentially converting lactate to pyruvate, and NAD+ to NADH^29^. The so-called “Warburg effect”, which was first described in cancer cells and later identified in immune cells, comprises a metabolic switch from oxidative metabolism towards glycolysis without a hypoxic environment, namely aerobic glycolysis preference, which is dependent on LDHA induction^30^. Overall, transcriptomic analysis and lactate determination of both microglia indicated an induction of glycolytic metabolism without hypoxic conditions.

**Figure 7.**
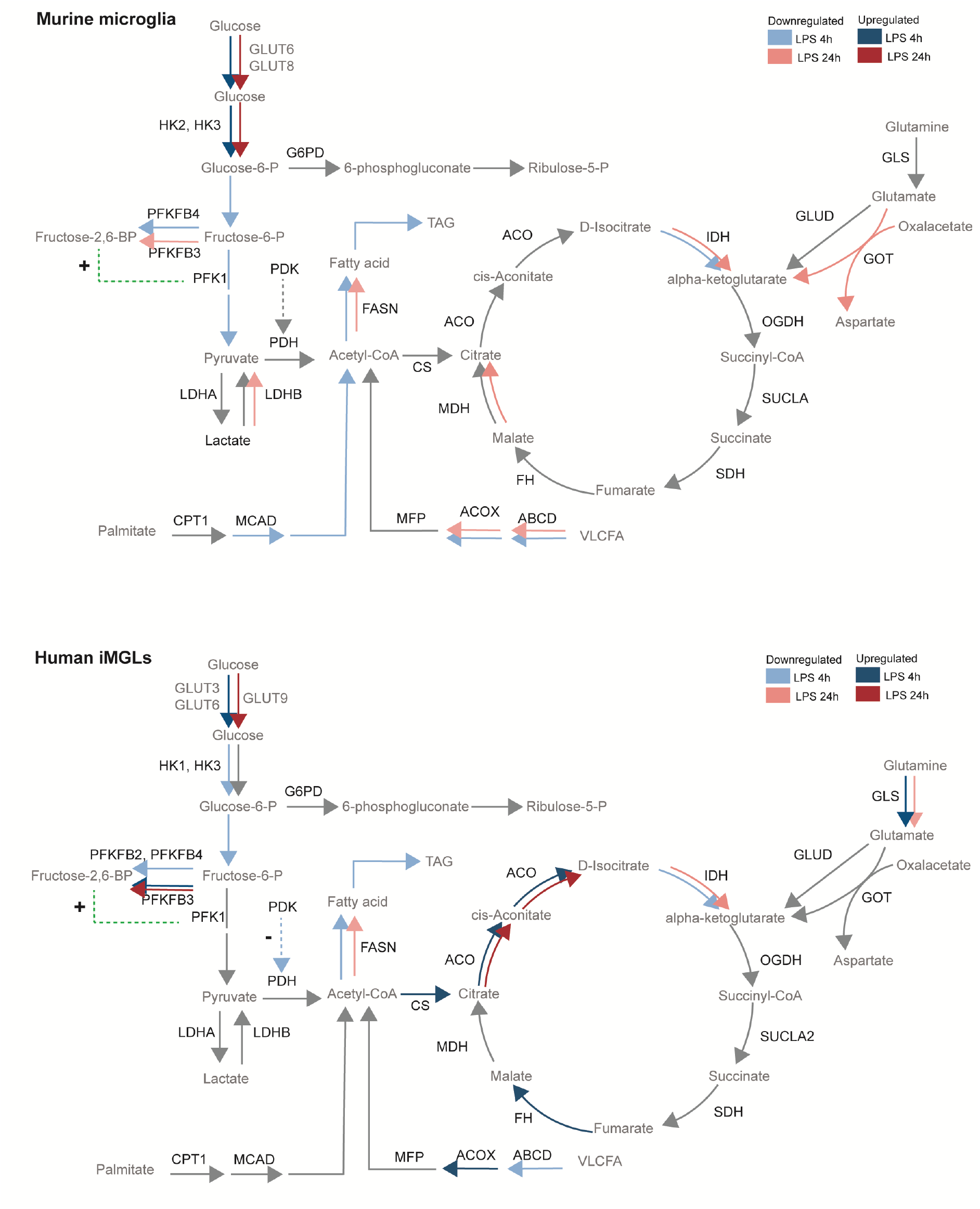
Similarities and divergences in mouse and human microglial metabolism. Upper panel: Upon LPS treatment mouse microglia upregulate genes that code for glucose transporters GLUT6 and GLUT8, HK2 and HK3 robustly at 4h and 24h with a shared decrease of downstream pathway components and downregulation of isocitrate dehydrogenase. A sustained downregulation of genes that encode key fatty acid import mediators, such as ATP-binding cassette transporter subfamily D (Abcd) and peroxisomal acyl-coenzyme A oxidase (Acox) is concomitant with a downregulation of fatty acid synthesis key enzyme fatty acid synthase (Fasn). Lower panel: Upon LPS treatment human iMGL cells robustly upregulate genes that code for glucose transporters GLUT3 and GLUT6 at 4h, and GLUT9 at 24h. Human iMGLs only upregulate hexokinases 1 and 3 at 4h, not at 24h. Downstream elements of the glycolytic pathway such as PFKFB3 are upregulated. An increase in Fructose-2,6 biphosphate promotes PFK1 activity. Within the TCA cycle, citrate synthase (CS) and aconitase (ACO) were upregulated both at 4h and 24h, while isocitrate dehydrogenases (IDH1 and IDH2) were downregulated both at 4h and 24h. Downregulation of genes that encode fatty acid import and catabolism, such as ABCD and ACOX occurs concomitantly with a sustained downregulation of FASN upon LPS. Grey lines denote unchanged gene expression. Dotted green lines denote promotion of enzyme activity. Dotted blue lines denote negative feedback on enzyme activity. This figure was created with BioRender.com.

The differentiated human iMGL cells resembled human adult and fetal microglia by both transcriptomic and functional analyses. Whole-transcriptome analysis of our iMGL cells and other cell types demonstrates a clear differentiation course of iPSCs to iHPCs and then microglia, clustering far from monocytes and dendritic cells. Traditionally, microglial differentiations strategies have been criticized and described to provide monocytic-like *in vitro* phenotypes, similar to cultured primary human microglia ^1^. As demonstrated by the study of Gosselin and colleagues ^2^, brain environment signals, such as TGF-β1 are necessary for the maintenance of the *in vivo* phenotype of microglia. Indeed, we have maturated microglial progenitors in the presence of IL-34, the tissue-specific ligand of CSF-1 receptor (CSF-1R) ^31^, and TGF-β1. Our experiments provide functional microglia-like cells that can be used to study mechanisms of disease and effects of brain microenvironment in the future.

The essential steps of glycolysis involving hexokinases, phosphofructokinase-1 were dysregulated in the transcriptomic datasets of both mouse and human microglia. When any fast (∼4h) transcriptomic dysregulation occurred, no changes in metabolism were observed. In contrast, long LPS stimulation (24h) elicited robust metabolic changes but subtle transcriptomic changes in all models. This discrepancy highlights the timeline between transcriptomic, translational, and functional responses in our model. These data, coupled to lactate measurements suggest that 4h LPS could represent a timepoint to assess transcriptomic induction of glycolytic genes, while functional responses may be detected later between 8h and 12h.

It is generally well-accepted that macrophages undergo metabolic reprogramming upon LPS treatment, a vital process for inflammatory responses^32–37^. Experiments with murine microglia and human microglial cell lines^38,39^ recapitulate key hallmarks of previous macrophage studies, such as the upregulation of glycolysis activity. Indeed, we observed an overall promotion of glycolysis, coupled to mitochondrial dysfunction in mouse microglia, supported by transcriptomic and functional assays. However, unlike mouse microglia, human microglia exhibited an increase of glycolysis without strict mitochondrial dysfunction. Our findings are in line with a recent study by Geric and colleagues^21^ which reported increased glycolytic flux upon LPS stimulation in mouse microglia with attenuation of oxidative metabolism. Here we report for the first time that LPS-treated human iMGL cells do not exhibit mitochondrial dysfunction coupled to glycolysis induction. We speculate that there is a species-specific mechanism that confers metabolic protection to human microglia. In particular, mechanisms that are linked to immune tolerance and a plethora of microglial phenotypes remain to be investigated.

Immune tolerance is a host-protective mechanism in which cells become unresponsive to subsequent stimulation and has been proposed to underlie pathophysiology of neurodegenerative diseases. LPS-induced maternal inflammation during late gestation was shown to affect microglial responsiveness during adulthood differentially between hippocampi and total brain microglia^40^, hinting tolerogenic mechanisms. It has been reported that tolerogenic murine microglia tend to normalize their glycolytic metabolism and increase their oxidative metabolism^13^, rendering them dysfunctional with poor phagocytosis capacity, chemotaxis, and inflammatory responses. Treatment with IFN-γ boosted glycolytic metabolism and reversed the defective microglial responses. However, human studies on the context of immune tolerance and metabolic reprogramming will be useful to assess whether boosting glycolysis in the chronic setting or inhibiting glycolysis in the acute setting will prove beneficial for therapy. A recent study reported that mature dendritic cells exhibit a metabolic signature consistent with upregulation of OXPHOS protein complexes, higher oxygen consumption ratio compared to immature dendritic cells^41^. Our data show that human iPSC-microglia exhibit these features. Whether human mature innate immune cells exhibit this metabolic signature remains to be further investigated.

Reports of human microglial phenotypes and subpopulations that are associated to disease hallmarks, aging or metabolic state have demonstrated that transcription of HKs or PFKs are differentially regulated in association with amyloid-beta responses – similarly to inflammatory microglia, and tau responses – similarly to homeostatic microglia^4,42^. Some studies have proposed glucose transporters^43^, HK2, and PFKs as therapeutic targets in neurodegenerative diseases with altered microglial functions aiming to attenuate glycolytic fluxes. However, we speculate that this will be beneficial only depending on the specific metabolic signatures that these microglia exhibit.

## Conclusion

In conclusion, we have shown a systematic comparison between human and mouse responses to the TLR4 agonist LPS. Despite the fact that human and mouse immune system share extensive similarities, as we observed here with the glycolytic upregulation in mouse microglia (*in vitro* and *in vivo*) and human iMGLs, many divergence pathways have been observed as well. For example, several enzymes such as Pfkl, Pfkp, Pfkb3, Pfkb4, Aco1, Mdh1 that are involved in glucose and TCA metabolism, are differentially regulated after LPS stimulation in mouse and human microglial cells. The increases in glycolytic activity are mainly attributed to hexokinase activity in mouse, and phosphofructokinase activity in human microglia. Considering that previous studies have not directly compared the energy metabolism between iMGLs and mouse microglia, this study presents the transcriptome profile and functional data on mouse and human microglia delineating their capacity to undergo metabolic reprogramming and confirming the relevance in translational research, which could provide a platform for targeting strategies for conditions associated with inflammation.

## MATERIALS AND METHODS

### Animals

C57BL/6 J mice from the central animal laboratory at University of Groningen were housed and handled in accordance to Dutch standards and guidelines (Protocol 171224-01-003). All experiments were approved by the University of Groningen Committee for Animal Experimentation.

### Primary Microglia Culture

Microglia cultures were prepared as previously described^44^. Briefly, brains were removed from 1-to 3-day-old C57Bl/6 pups, minced, dissociated for 25 min in 0.25% Trypsin, 1xDNAse, and then cells were cultured in Dulbecco’s modified Eagle containing (DMEM, Gibco, 42430025), supplemented with 10% fetal bovine serum (FBS, Hyclone, SV30160),1 mM Sodium Pyruvate (ThermoFisher, 15070063), and 100 U/mL pen/strep (ThermoFisher, 11360070) in the incubator at 37°C and 5% CO_2_. After 2 days of *in vitro* cultivation, the growth medium was completely replaced by fresh medium. After 10–14 days, flasks were mechanically shaken for 60 min, 150 rpm to yield microglia in the supernatant, which were sub-cultured into uncoated well plates according to the experiment. They were kept in 50% astrocyte conditioned medium and 50% fresh DMEM supplemented with 10% FBS, 100 U/mL pen/strep. For all experiments, primary microglial cells were used only for the first and second passage.

### Animals (*In vivo* treatment)

The transcriptome analysis of the TCA/glycolytic genes of *in vivo* microglia was performed using our recently published data (Zhang et. al, 2022)^20,45^.

### Maintenance human induced-Pluripotent Stem Cells

The induced pluripotent stem cell (iPSC) cell line Ctr8.2 (UEF-2B) was obtained from Virtanen Institute for Molecular Sciences-University of Eastern Finland^46^. Another iPSC line purchased from Gibco was also used for the functional assays. The iPSC colonies were cultured and expanded onto Matrigel (Corning, 354277) in 6 well plates and mTeSR8 medium (Gibco, A1516901) supplemented with 100 U/mL penicillin/Streptomycin. Once the cells were confluent, iPSC colonies were passaged every 3-4 days using enzymatic detachment with EDTA 0,5 mM for 5 min and re-plated in mTeSR8 medium with RevitaCell (Gibco, A2644501). The pluripotency of iPSCs was tested regularly by staining for the pluripotency-marker OCT4. In addition, the cells were examined regularly for the presence of mycoplasma.

### Microglia differentiation protocol from iPSCs

Microglia differentiation from iPSCs was based on previously described protocols^22^. Day 0 of differentiation was started when the cells reached 70 to 80% confluency. Cells were detached with accutase (Sigma Aldrich, A6964) and incubated at 37°C for 5 minutes. Detached cells were transferred to a Falcon tube and centrifuged for 5 minutes 300rcf. Cells were resuspended in differentiation medium mTers1 supplemented with fresh cytokines with 50 ng/mL BMP-4, 50 ng/mL VEGF, 20 ng/mL SCF (differentiation medium 1) and 10 μM Rock Inhibitor. 10.000-15.000 cells per well were seeded in a 96-well U-bottom ultra-low adherence plate (Corning, 7007) and the plate was centrifuged at 100 rcf for 5 minutes to ensure clustering of the cells at the bottom of the well. The plate was incubated at 37°C, 5% CO_2_.

From day 1 to day 3, 75% of medium was changed using differentiation medium 1. At day 4, all the embryoid bodies (EBs) were collected and medium was removed. The EBs were resuspended in differentiation medium 2 (XVIVO 15 medium + 2mM Glutamax, 100 U/mL Pen-Strep and 0.055 mM 2-mercaptoethanol with 50 ng/mL SCF, 50 ng/mL M-CSF, 50 ng/mL IL3, 50 ng/mL FLT3 and 5 ng/mL TPO). Approximately 20 EBs were plated per well of a 6-well plate. At day 8, medium was changed by removing all the medium in the wells and adding new differentiation medium 2. At day 11, the old medium was removed and the EBs were resuspended in 20 mL of differentiation medium 3 (XVIVO 15 + 2mM Glutamax, 100 U/mL Pen-Strep and 0.055 mM 2--mercaptoethanol with 50 ng/mL FLT-3, 50 ng/mL M-CSF and 25 ng/mL of GM-CSF). At day 18, the supernatant was collected and passed through a 37 μm cell strainer and spun down at 300 rcf for 5 min. Microglial progenitors were resuspended in maturation medium (Advanced DMEM/F12 supplemented with 5 μg/mL N-acetylcysteine, 400 μM 1-Thioglycerol, 1 μg/mL heparan sulfate, 1% Glutamax, 1% NEAA, 1% Pen-Strep, 2% B27, and 0.5% N2. Growth factors: 100 ng/mL interleukin-34, 25 ng/mL M-CSF, 25 ng/mL CX3CL1, 25 ng/mL TGF-β-1 and 50 ng/mL CD200) and used for experiments and termed throughout the manuscript as iPSC-derived microglia-like (iMGL) cells.

### IncuCyte Phagocytic Assay

For the Phagocytic assay, 15,000 cells were seeded in a 96-well flat bottom plate in DMEM (4.5 g/L) or maturation medium, for mouse or iMGL, respectively. The cells were incubated for 24hr at 37°C and 5% CO_2_. 4hr before treatment, the plate was placed inside the IncuCyte S3 to obtain a blank measurement of the cells. The cells were treated with Cytochalasin D (Sigma Aldrich, C8273, 10μM) in microglia medium with pHrodo red *E*.*coli* bioparticles 1ug/0.1ml (Thermo Fischer, p35361) and monitored for a minimum of 24hr. The data was obtained using the IncuCyte S3.

### Immunofluorescent staining

Cells were fixed using paraformaldehyde (PFA) 4% for 20 min and permeabilized using Triton X-100 0.1% for 10 min. Unspecific binding was blocked by incubation with 5% BSA for 1hr at room temperature. Incubation with primary antibody against IBA1, OCT4, was conducted overnight at 4°C. Secondary anti-rabbit antibodies coupled to Alexa Fluor 488 (Invitrogen, A11034) or 568, (Invitrogen, A10037) were incubated at room temperature. Nuclei were counterstained with DAPI. Images were acquired using a Leica DFC3000 G camera and acquired using Leica Las 4.3 software.

### Real-time cell impedance measurements

Morphological changes in mouse and iMGL cells were measured using a label-free, real-time cell impedance-based system; xCELLigence® RTCA MP system (ACEA BIO). 15,000 cells/well were seeded in a 96-well E-plate (Agilent, 5232368001)^47^ containing gold microelectrodes fused to the bottom surface of the well plate. The impedance of the electron flow caused by cell attachment to the well (cell index) was measured every 30 minutes. The cells were seeded one day before treatment and the measurement of the cell index was performed 24hr following LPS treatment. The cell index was normalized to 1, before the LPS stimulation.

### Mitochondrial respiration measurement

Measurement of oxygen consumption rate (OCR) and extracellular acidification rate (ECAR) was performed using XFe extracellular flux analyzer (Seahorse Bioscience, Billerica, MA) and mitochondrial stress test kit. 50,000-70,000 cells/well of human or mouse microglia cells were seeded in XFe96 cell culture microplate. 4 or 24hr following addition of LPS stimulation, the medium was replaced by Seahorse XF base medium (Agilent, 102353-100) supplemented with 10 mM glucose, 1 mM sodium pyruvate and 2 mM glutamine and incubated for 1 hr in CO_2_ free incubator at 37 °C. The following substances were injected in order; port A:

Oligomycin (2.5 μM), port B: FCCP (0.5 μM) and port C: rotenone (0.5 μM) and antimycin A (0.5 μM). Three baseline OCR measurements (3 min mix, 0 min delay, 3 min measure = 3/0/3) were recorded, followed by assessment of mitochondrial metabolism by injection of oligomycin (3/0/3), FCCP (3/0/3), and a combination of rotenone and antimycin A (3/0/3). The following parameters were deduced: basal respiration (OCR values, used to provide ATP under baseline conditions), ATP-linked respiration (following oligomycin injection, a reduction in OCR values represents the part of basal respiration used to produce ATP), maximal respiration (the maximal OCR values following FCCP injection)^48,49^. Basal glycolysis was calculated from the subtraction between the maximum measurement before oligomycin injection and the last measurement before glucose injection. Glycolytic capacity was calculated by subtracting the maximum measurement after oligomycin injection and the last measurement before glucose injection. Glycolytic reserve was calculated by subtracting glycolysis capacity and basal glycolysis and it is represented as a percentage. At least 3 independent experiments were performed. Two-way ANOVA test was used to determine statistical significance for the different mitochondrial parameters.

### Lactate measurement

Cells were seeded in a density of 15,000 cells/well and treated with LPS 250 ng/ml in DMEM or maturation medium, for mouse or iMGL, respectively. Lactate release was measured after 4-8 h of treatments. Medium from microglia cells was collected and diluted 3 times in demineralized water. A calibration curve of eight lactate standards ranging from 0 to 1.2 mM was prepared for quantification purposes. Subsequently, lactate was measured in a 96-well plate using 20μl medium sample or lactate standard mixed with 225 μl reaction mixture (0.44 M Glycine/ 0.38 M Hydrazine [pH 9.0], 2.8 mM NAD) and 5 units L-lactic dehydrogenase (EC 1.1.1.27) followed by absorbance determination at 340 nm using the 120 SynergyTM H4. All chemicals were purchased from Merck Millipore. Background absorbance of the blank control (0.0 mM lactate standard) was subtracted from all sample readings and medium samples were corrected for dilution. Medium lactate concentrations were determined based on linear regression of the standard curve.

### Treatment and RNA isolation

Mouse microglia was treated with LPS (250ng/m) for 4 or 24hr. The control cells were collected following 24h media change. Isolation of RNA was performed using the Nucleospin® RNA isolation Kit (Macherey-Nagel). The samples were stored at -80°C until they were sent for transcriptomic analysis.

### Library construction and sequencing

The quality of the samples, the construction and the sequencing of the RNA-libraries were performed by GenomeScan Bv. Data obtained from the transcriptomic analysis: quality control data, raw count files, FPKMs, RPMs and CPMs.

### RNA-seq analysis

Differential gene expression analysis was performed with the DESeq2 package^50^ For human microglia RNA-seq, batch effect differences were controlled by including the collection number in the design formula. Several comparisons were made, for all we used an absolute log fold change >1.5 and an FDR-adjusted p-value <0.05 to define a differentially expressed gene (DEG). Gene set enrichment analysis (GSEA) for individual comparisons was performed using Metascape. For visual comparison of the changes of Glycolytic and TCA pathways upon LPS treatments between human and mouse microglia, the Log2 Fold Changes of 1-to-1 ortholog genes were plotted as scatter plots. One to one orthologs between mouse and human were obtained from Biomart (https://www.ensembl.org/). Visualizations were made with the packages ‘ggplot2’ and ‘pheatmap’.

### Statistical analysis and visualization

Statistical analysis was performed using a paired student’s T test or ANOVA and Tukey’s test for post hoc multiple comparisons of the parametric data in between-group analyses. Non-parametric data were evaluated using Kruskal-Wallis test. Data were analyzed using GraphPad Prism software (version 9.0, GraphPad Software Inc., La Jolla, CA, USA), expressed as mean ± SD for transcriptomic data and SEM for the rest experiments. DESeq2 analysis used Wald test to determine significance. Graphs were visualized using Excel and GraphPad Prism version 9. P values indicating statistical significance differences between mean values are defined as follows: ^*^*p*< 0.05, ^**^*p* < 0.01, ^***^*p* < 0.001.

## Acknowledgments

AMD is a recipient of a Rosalind Franklin Fellowship co-funded by the European Union and the University of Groningen. This work was partially funded and supported by Parkinson Fonds. We acknowledge support from Renzo Mancuso lab, Nicola Fattorelli and Ana Martinez Muriana for the development of iMGLs protocol.

## Author Contribution

A.M.S.G., A.M.G: Conceptualization, Methodology, Investigation, Visualization, Writing -Original Draft. M.T.L., A.O., T.C., and J.H.: Methodology, Investigation, Visualization, A.K., B.E., E.B., B.M.B., S.L. and J.K.: Resources, Methodology, Writing - Review & Editing, Supervision. AMD: Conceptualization, Methodology, Resources, Writing - Original Draft, Review & Editing, Supervision, Funding acquisition. All authors have read and agreed to the published version of the manuscript.

